# Murine Progeria Model Exhibits Delayed Fracture Healing with Dysregulated Local Immune Response

**DOI:** 10.1101/2024.05.29.596277

**Authors:** Victoria R. Duke, Marc J. Philippon, Dane R.G. Lind, Herbert Kasler, Kohei Yamaura, Matt Huard, Molly Czachor, Justin Hollenbeck, Justin Brown, Alex Garcia, Naomasa Fukase, Ralph S. Marcucio, Anna-Laura Nelson, William S. Hambright, Dustin M. Snapper, Johnny Huard, Chelsea S. Bahney

## Abstract

**Background:** Bone fracture is one of the most globally prevalent injuries, with an estimated 189 million bone fractures occurring annually. Delayed union or nonunion occurs in up to 15% of fractures and involves the interruption or complete failure of bone continuity following fracture. Preclinical testing is essential to support the translation of novel strategies to promote improved fracture repair treatment, but there is a paucity of small animal models that recapitulate clinical attributes associated with delayed fracture healing. This study explores whether the *Zmpste24*^-/-^ (Z24^-/-^) knockout mouse model of Hutchinson-Gilford progeria syndrome presents with delayed fracture healing. Leveraging the previously characterized Z24^-/-^ phenotype of genomic instability, epigenetic changes, and fragility, we hypothesize that these underlying alterations will lead to significantly delayed fracture healing relative to age-matched wild type (WT) controls.

**Methods:** WT and Z24^-/-^ mice received intramedullary fixed tibia fractures at ∼12 weeks of age. Mice were sacrificed throughout the time course of repair for the collection of organs that would provide information regarding the local (fracture callus, bone marrow, inguinal lymph nodes) versus peripheral (peripheral blood, contralateral tibia, abdominal organs) tissue microenvironments. Analyses of these specimens include histomorphometry, μCT, mechanical strength testing, protein quantification, gene expression analysis, flow cytometry for cellular senescence, and immunophenotyping.

**Results:** Z24^-/-^ mice demonstrated a significantly delayed rate of healing compared to WT mice with consistently smaller fracture calli containing higher proportion of cartilage and less bone after injury. Cellular senescence and pro-inflammatory cytokines were elevated in the Z24^-/-^ mice before and after fracture. These mice further presented with a dysregulated immune system, exhibiting generally decreased lymphopoiesis and increased myelopoiesis locally in the bone marrow, with more naïve and less memory T cell but greater myeloid activation systemically in the peripheral blood. Surprisingly, the ipsilateral lymph nodes had increased T cell activation and other pro-inflammatory NK and myeloid cells, suggesting that elevated myeloid abundance and activation contributes to an injury-specific hyperactivation of T cells.

**Conclusion:** Taken together, these data establish the Z24^-/-^ progeria mouse as a model of delayed fracture healing that exhibits decreased bone in the fracture callus, with weaker overall bone quality, immune dysregulation, and increased cellular senescence. Based on this mechanism for delayed healing, we propose this Z24^-/-^ progeria mouse model could be useful in testing novel therapeutics that could address delayed healing.

**The Translational Potential of this Article:** This study employs a novel animal model for delayed fracture healing that researchers can use to screen fracture healing therapeutics to address the globally prevalent issue of aberrant fracture healing.

## INTRODUCTION

In 2019 the Lancet Global Burden of Disease estimated 189 million fractures occurred annually, an increase of 33.4% since 1990 [1]. While many fractures heal completely without incident, there remains a high prevalence of fractures with impaired healing, with the highest rates found in femoral (13.6%) and tibial (11.7%) shaft fractures [2, 3]. While there remains no single consensus for the clinical definitions of delayed and nonunion, generally, delayed union may be considered for long bone fractures with limited evidence of healing between 3-6 months and nonunion is diagnosed following the failure of a fracture to heal at 9 months [4, 5]. Delayed union and nonunion fractures commonly require several surgical procedures to achieve healing significantly extending the treatment period and contributing to lengthy patient disability, persistent pain, and increased healthcare costs [2, 6]. The incidence rate for osteoporotic/osteopenic fractures is significantly increased in the elderly population and contributes to morbidity and mortality in aged individuals [7]. Further, the underlying physiological landscape characterized by chronic low-grade inflammation is known to disrupt proper bone healing [8]. Given that in the next 30 years the global population of individuals over the age of 65 is projected to double, reaching 1.5 billion by the year 2050 [9], it is critical to develop therapeutics that promote the regeneration of bone after injury to meet the medical challenges faced by an aging population.

One barrier to discovering novel regenerative therapeutics to address delayed healing in the aging population is an unmet need for comprehensive and practical preclinical models. While a variety of preclinical models have been developed, many are associated with limitations as they typically model only a specific cause of delayed healing. The most prevalent models of delayed union in rodents are physically induced via segmental defects, modified fixation stiffness, or additional soft tissue, vascular, or periosteal injury [5, 10–12]. These models are useful for screening potential therapeutics that address significant trauma scenarios and investigate how mechanical factors such as weight-bearing and fixation properties influence regeneration. However, mechanically induced nonunion is often performed on healthy mice and thus does not incorporate the perturbed immunological state that drives delayed union in aging populations [8].

Naturally aged mice have been effectively studied and used to reveal important biological mechanisms underlying delayed fracture healing, most prominently immune and osteoanabolic dysregulation [13–15]. However, the NIH standard for age-related research in mice is 24 months, a constraint that creates a substantial time and monetary burden for effectively screening therapies associated with age-related decline in bone healing. Alternatively, preclinical models that induce diabetes mellitus or rheumatoid arthritis have proven useful in modeling impaired bone union in the presence of chronic inflammation [16–18]. While these models are clinically relevant to the targeted disease, they often exhibit severely dysregulated immune responses that are absent from the more subtle pathology of fracture healing in the general aged population. Additionally, patients at risk of delayed union commonly present with a frailty phenotype established to negatively influence bone regeneration, including osteoporosis and sarcopenia, that existing fracture models do not capture. As such, there is an unmet need to develop more practical preclinical models of delayed fracture healing in the aged population.

Hutchinson-Gilford progeria syndrome (HGPS) is a systemic disease of accelerated aging caused by a mutation in the nuclear protein Lamin A [19]. The disrupted nuclear lamin structure alters gene expression and genomic stability, resulting in biological changes observed in natural aging, including DNA damage, cellular senescence, cytoskeletal stiffness, and stem cell exhaustion [20–23]. Of the various murine models of HGPS established to research age-related diseases, genetic deletion of *Zmpste24 –* a metalloprotease essential for processing prelamin A into mature lamin A – recapitulates the premature aging phenotype with accompanying musculoskeletal deficits reflective of frail individuals [24–26]. Within 3-4 months of age, the *Zmpste24-/-* (Z24^-/-^) mouse model exhibits musculoskeletal deficiencies, including reduced myogenic and osteogenic stem cell proliferation and differentiation, bone density loss, sarcopenia, weight loss, osteoporosis and osteoarthritis [20, 27]. Further, frail individuals have also been shown to have reduced circulating osteoprogenitors and reduced lamin A, providing evidence that the Z24^-/-^ mouse model may have translational relevance to delayed bone regeneration in the context of aged fractures [28].

Thus, the goal of this study was to leverage the Z24^-/-^ progeria mouse as a model of delayed fracture healing in the tibia, evaluating both the fracture and immune phenotype following injury. We hypothesized that Z24^-/-^ mice would present with delayed fracture healing when compared to age-matched wild-type (WT) controls and exhibit both increased senescent cell burden and a dysregulated immune response. Because the complex mechanics and biology of delayed and nonunion fractures remain poorly understood, management of these injuries poses a significant challenge to clinicians in deciding to either wait for healing to occur or intervene with either surgical or non-surgical methods. The fracture healing phenotype of the Z24^-/-^ mouse represents a new preclinical translational tool that is more feasible than natural aging to test efficacy and mechanism of action for novel therapeutic strategies designed to accelerate bone healing in aged individuals.

## MATERIALS AND METHODS

### Animal model

A Material Transfer Agreement (MTA) was executed with the Universidad de Oviedo and Dr. Carlos Lopez Otin for the use of the *Zmpste24* mice (Recipient Scientists: Drs Johnny Huard and Chelsea Bahney). Z24^-/-^ progeria mice were generated as previously described through a homozygous deletion of *Zmpste24* on the C57Bl6/J background [20]. For the present study, Z24^-/-^ mice were created by crossing two heterozygous Z24^-/+^ mice and congenic Z24^+/+^ littermates were used as WT controls. Genotyping was performed using a three-primer mix (**Table 1**). WT mice were identified by a DNA band at 520 base pairs and Z24^-/-^ mice were identified by a DNA band at 303 base pairs. All procedures received approval from the Colorado State University Institutional Animal Care and Use Committee (IACUC) and followed NIH guidelines for ethical treatment of animals.

**Table 1.**
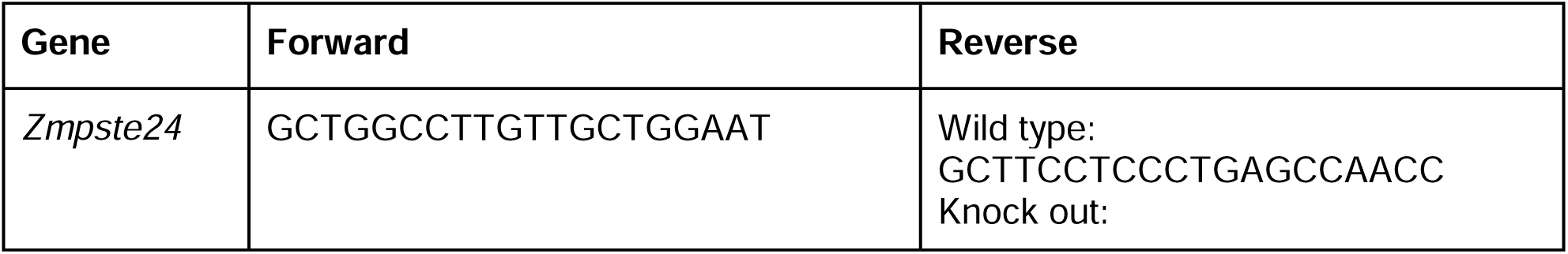
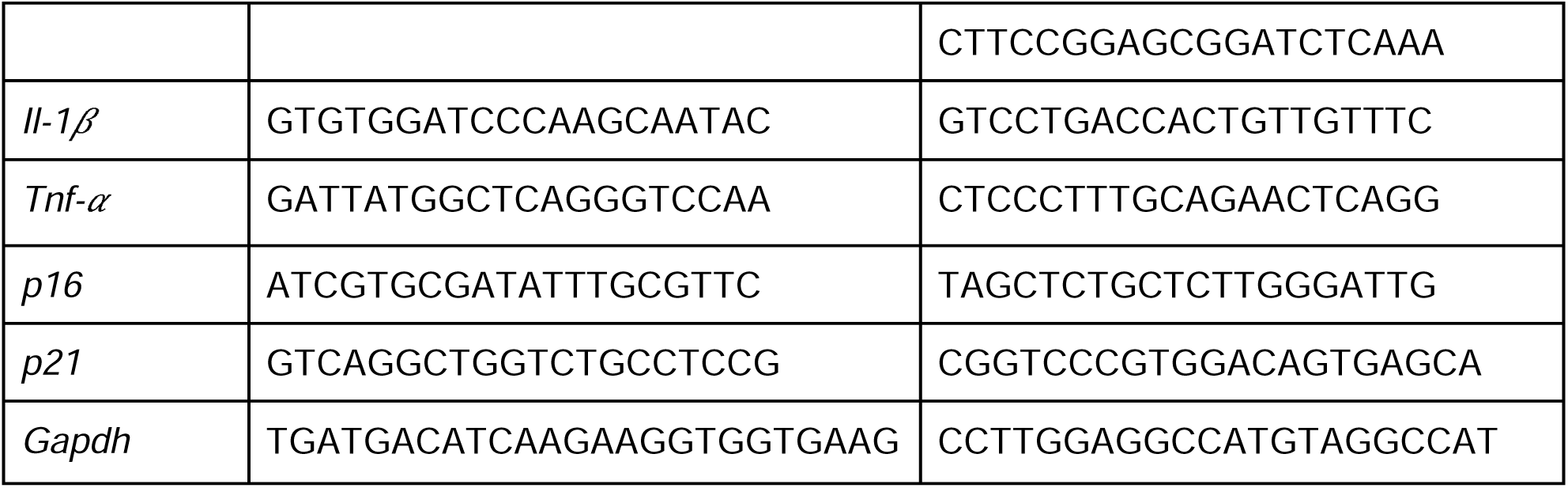
Mouse Primer Sequences

### Murine tibial fracture and stabilization model

Z24^-/-^ and WT mice (10-14 weeks) were anesthetized using isoflurane inhalation. The right leg was prepared for surgery by shaving the surgical site, then disinfecting with three rounds of 70% alcohol wipes followed by chlorhexidine surgical scrub solution. Sustained release buprenorphine (0.6-1.0 mg/kg) was then administered and mouse eyes were lubricated using artificial tears ointment. All surgeries were performed on a heated operating table using aseptic technique. To create the fracture, a small incision was made along the tibia, and a 23-gauge needle was used to form a hole at the top of the tibial plateau. A sterilized intramedullary pin was inserted through the hole spanning from the tibial plateau, through the tibial cavity, and into the distal tibia. A Dremel was used to create two small holes in the mid-shaft of the tibia and pressure was applied to both the proximal and distal ends to generate a full-thickness tibial fracture as previously described [29]. The incision was closed with 5-0 Biosyn Sutures (Covidien, 5687) and one surgical skin staple. Bupivacaine hydrochloride (NovaPlus, RL7562) was applied topically for post-operative pain management. Mice were socially housed and allowed to ambulate freely with 72 hours of post-operative monitoring. DietGel® nutritional supplements (ClearH_2_O®, 72085022) were provided to all mice 14 days prior to and post-surgery until euthanasia to maintain body weight. Animals were euthanized 3-, 9-, 14-, or 21 days post-fracture by carbon dioxide (CO_2_) asphyxiation and tissues harvested for analysis (**Figure 1**).

**Figure 1.**
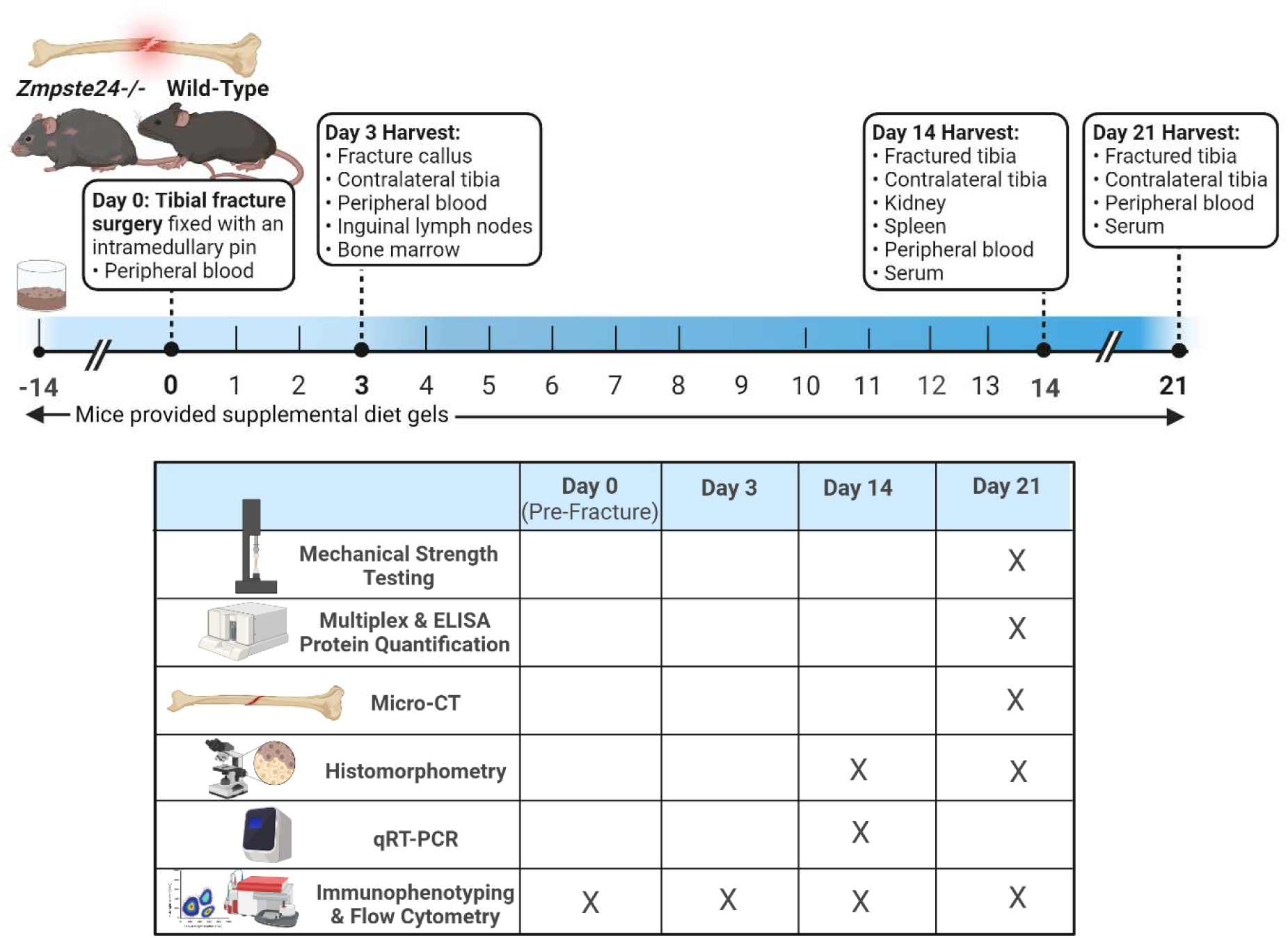
**Schematic representation of experimental design**. The experiment duration varied from 3 to 21 days, depending on the tissues harvested. At day zero (mice 10-14 weeks of age), surgery was performed to create a tibial fracture that was stabilized with an intramedullary pin. Mice were allowed to ambulate freely until euthanized at 3-, 14-, or 21 days post-fracture by CO_2_ asphyxiation for tissue collection. Figure created with BioRender.com.

### Histology and histomorphometry

Fractured tibiae were dissected 14 and 21 days post-fracture and fixed in 4 % paraformaldehyde (Santa Cruz, SC281692) for 24 hours at 4 °C. Bones were then decalcified by rocking in 19 % ethylenediaminetetraacetic acid (Biorad, 1610729) for 14 days at 4 °C with solution changes every other day. Decalcified tibiae were paraffin embedded and whole samples were serially sectioned into 8 µm sections with 3 sections per slide. Samples were stained with Hall Brundt’s Quadruple staining protocol to visualize osseous (red) and cartilaginous (blue) tissue within the fracture callus as previously described [30].

For histomorphometric quantification of the tissue composition in the fracture callus, the first section of every tenth slide was imaged using a Nikon Eclipse Ni-U microscope with Nikon NIS Basic Research Elements Software (Nikon Instruments Inc., version 4.30) captured at 2X magnification. The fracture callus region of interest for each image was isolated using the “Lasso” tool on Adobe Photoshop (version 23.0.1) to remove non-callus tissue from each image. The area of each callus tissue (cartilage, bone, and background) was then quantified using the Trainable Weka Segmentation add-on in Fiji ImageJ (version 1.51.23; NIH), as previously described [31]. The total volume of each tissue type in the fracture callus was approximated by multiplying the tissue area on each slide by the uniform spacing between each slide over the entire length of the fracture callus. The volumes of each tissue type are expressed as percentages of the total fracture callus volume per mouse.

### Micro-computed tomography (**μ**CT) analysis

Tibiae were collected 21 days post-fracture after euthanasia to assess bone formation from the injured site through morphometric analysis performed with *ex vivo* μCT scanner (Scanco Medical AG, CH-8306, Bruttisellen, Switzerland) with the following parameters: 70kVp; filter: 0.5 mm AL; BH:1200 mg HA/ccm; scaling 4096; voxel size: 15.6 µm. Three-dimensional images were reconstructed using Scanco Medical Software. Quantification of total volume (TV), bone volume (BV), bone mineral density (BMD), trabecular number (Tb.N), and trabecular thickness (Tb.Th) were performed in the defect region.

### Mechanical Strength Testing

Fractured and contralateral tibiae were collected 21 days post-fracture, wrapped in phosphate-buffered saline (PBS)-soaked paper towels, and stored at 4°C until testing. The proximal and distal ends of the tibiae were potted in a cylindrical mold using polymethyl methacrylate (Fricke Dental) such that the longitudinal axis of the proximal and distal cylindrical molds was aligned to the anatomical axis of the tibia. Torsional testing to failure was performed in a blinded fashion. Using custom fixtures, each specimen was mounted to an Axial Torsion Test Machine (Test Resources, 131AT) instrumented with a 2 Newton-meter (N-m) capacity torque load cell. Torsional load was applied at 1 degree/ second until failure occurred. Failure torque (N-m) and torsional stiffness (N-m/degree) were measured. Failure torque was defined as the maximum force that the bone sustained before fracture, and torsional stiffness was calculated as the slope of the linear portion of the torque-rotation curve.

### Gene Expression Analysis

Fractured tibiae (day 9) and kidney (day 14) were harvested and stored in RNAlater (Invitrogen, AM7020) at -20 °C until RNA isolation. Fracture calli were carefully dissected from the tibiae/surrounding muscle and all organs were homogenized in Trizol reagent (Invitrogen, 15596026) on ice. Total RNA was extracted from samples according to the manufacturer’s protocol. Reverse transcription of RNA into cDNA was performed via the manufacturer’s protocol using qScript cDNA synthesis kit (VWR, 95048100). Quantitative RT-PCR (qRT-PCR) using PerfeCTa SYBR Green FastMix (VWR, 101414280) was run on the Step-One Plus Real-Time PCR system (Applied Biosystems,4376598). Relative gene expression of interleukin 1), tumor necrosis factor-alpha (*Tnf-α)*, and cell-cycle regulator proteins *p16* and *p21* was calculated by normalizing results to the housekeeping gene (*Gapdh*, ΔRelative gene expression was calculated using the 2^−ΔCT^ method.

### Senescence quantification in peripheral blood mononuclear cells

Peripheral blood was collected via tail vein pre-fracture and cardiac puncture 14 days post- fracture using a syringe coated with anticoagulant citrate-dextrose solution A (ACD-A; McKesson, 353996). Peripheral blood mononuclear cells (PBMCs) were isolated using pluriMate® centrifugation tubes pre-filled with PBMC Spin Medium® (pluriSelect, 440920215) and control rate frozen in CryoStor® (StemCell Technologies, 7930) until analysis. To quantify cellular senescence, C_12_FDG was used to label cells demonstrating elevated senescence-galactosidase activity [32, 33]. PBMC samples were briefly thawed and stained with 500 nM bafilomycin for false positive reduction (Cell Signaling Technology, 54645), 30 µM C_12_FDG (Abcam, ab273642), and DR to identify dead cells (NOVUS Biologicals, NBP2- 81126). Samples were run on a Guava® easyCyte™ HT (Guava®) flow cytometer and analyzed using InCyte software (GauvaSoft 3.3).

### Plasma protein quantification

Whole blood was collected (21 days post-fracture) and plasma was isolated using ACD-A coated tubes, followed by centrifugation at 1500 x g for 10 minutes. Supernatant was collected and stored at -80°C until analysis. Multiplexed chemokine and cytokine analysis was performed using Milliplex Mouse Cytokine/Chemokine Magnetic Bead Panel (EMD Millipore, MCYTOMAG70K10) according to the manufacturer’s protocol and read on a Luminex® 100/200 platform using xPonent® software (EMD Millipore Corp, LX200-XPON-RUO, version 4.2). All standards and samples were run in duplicate and analyte concentrations were calculated using Belysa® Immunoassay Curve Fitting Software (EMD Millipore, 40-122, version 3.4). Standard ELISA kits were used to quantify growth differentiation factor 15 (GDF-15) (R&D systems, MGD150) and sclerostin (SOST) (R&D systems, MSST00) protein levels following the manufacturer’s protocol. All standards and samples were run in duplicate using the Infinite® M plex microplate reader (Tecan, 30190085).

### Immunophenotyping

Inguinal lymph nodes, peripheral blood via the tail vein, and tibia and femur bone marrow were harvested between 0 and 14 days post-fracture (**Figure 1**). These samples were analyzed using a 31-color panel spectral flow panel for peripheral blood and lymph nodes and a 30-color panel spectral flow panel for bone marrow (**Supplementary** Figure 1). Single-cell suspensions were prepared from mouse tissues as follows: inguinal lymph nodes were placed in 6-well or 12-well plates, in 5 mL or 1 mL of ice-cold RPMI/10% FBS and crushed with a syringe plunger to dissociate the cells. To remove the stroma, both suspensions were then filtered through a 40µm nylon mesh (Genesee, 25-375) and pelleted at 600 x g for 6 minutes. Supernatants were discarded, and the cells were taken up in 3 mL or 1 mL of ACK lysis buffer as previously described [34] and incubated for 3 minutes at room temperature (RT). Afterward, 10 times the buffer volume of cold PBS was added and the cells were again pelleted as above. For peripheral blood, approximately 50 µL was collected from the tail vein into 1.4 mL of cold PBS/2 mM EDTA in microcentrifuge tubes, then pelleted for 4 minutes at 600 x g. After removal of the supernatant, the pellets were taken up in 350 µL ACK lysis buffer and incubated for 3 minutes at RT. Afterward, 1.2 mL of cold PBS was added, and the cells were again pelleted. For bone marrow, tibias and femurs were dissected from whole legs and crushed using a syringe plunger in 5 mL RPMI/10% FBS. After vigorous resuspension to dissociate the bone marrow, the suspension was filtered through a 40 µm nylon mesh (Genesee, 25-375), pelleted at 600 x g for 6 minutes, and lysed with ACK buffer, followed by dilution with 10 volumes of PBS and pelleting of the cells.

For each single-cell suspension, samples were then suspended in a minimal volume of PBS and counted. 0.5-2.0 x 10^6^ cells were transferred to 1 mL of RPMI/10% fetal bovine serum (FBS), pre-warmed to 37 °C in microcentrifuge tubes. Bafilomycin A1 (ThermoFisher Scientific, 328120001) was then added at 500 nM in DMSO, and the cells were incubated at 37 °C for 45 minutes. After this time, C_12_FDG (Invitrogen, D2893) was added to 30 µM final concentration and the cells were incubated at 37 °C for another 45 minutes. At the end of the C_12_FDG labeling, the cells were pelleted as above, washed once with 1 mL of cold PBS, taken up in 120 mL of PBS/2mM EDTA with 2X fixable Live/Dead Blue stain (Invitrogen, L23105), transferred to 96-well V-bottom plates, and incubated on ice in the dark for 15 minutes. After live/dead staining, the antibody cocktails were added neat in the noted volumes (**Supplementary Table1**) along with sufficient FBS to bring the final concentration to 2%, and the samples were incubated for another 30 minutes on ice in the dark. The samples were then diluted with 150 µL of PBS/2mM EDTA/2% FBS, pelleted at 600 x g for 5 minutes, and resuspended in 150 µL of PBS/2mM EDTA/2% FBS for acquisition. Sample acquisition was performed on a 5-laser Cytek Aurora spectral flow cytometer (Cytek Biosciences), and spectral unmixing was done using SpectroFlo software (Cytek). Subsequent manual correction of unmixing and gating was done using FlowJo software (BD Biosciences, version 10.1) to identify unique cell populations.

### Statistical Analysis

Statistical analyses were performed using GraphPad Prism (Version 9.4). The non-parametric Mann-Whitney U test was used to determine significance between two groups. Flow immunophenotyping data was analyzed using a two-tailed T-test. Comparisons among multiple groups were performed using two-way ANOVA followed by Tukey’s post-hoc testing, excluding time course data, which was analyzed using a mixed-effects model followed by Tukey’s post-hoc testing. All data are presented as either the mean ± standard deviation or the median ± standard error of mean from at least four independent experiments with biological replicate specified in each figure. P < 0.05 was used to indicate statistical significance.

## RESULTS

### Z24^-/-^ mice exhibit delayed fracture healing and decreased torsional bone strength in unfractured tibia compared to age-matched WT mice

At both 14 and 21 days post-fracture, fracture calli from Z24^-/-^ mice grossly displayed more cartilage (blue) and less bone (red) tissue compared to that of age-matched WT mice (**Figure 2A, B, D, E**). Quantitative histomorphometry revealed 37.1% lower bone volume (p = 0.029) and 107.0% higher cartilage tissue volume (p = 0.0286) in Z24^-/-^ mice compared to WT mice 14 days post-fracture (**Figure 2G**). This trend persisted for 21 days post-fracture with Z24^-/-^ mice demonstrating a 33.7% reduction in bone volume (p = 0.016) and a 379.7% (p = 0.0159) increase in cartilage volume (**Figure 2H**). Likewise, at 21 days post-fracture, μCT analysis indicated that both TV (p = 0.013) and BV (0.0047) were lower in Z24^-/-^ mice. BMD within the fracture callus also trended lower in Z24^-/-^ mice (p = 0.2284, **Figure 2I**). The mechanical integrity of the unfractured, contralateral tibiae of Z24^-/-^ mice demonstrated average failure torque and torsional stiffness that were 48.0% (p = 0.0087, **Fig. 2J)** and 31.9% (p = 0.0303, **Figure 2K**) lower than WT mice, respectively.

**Figure 2.**
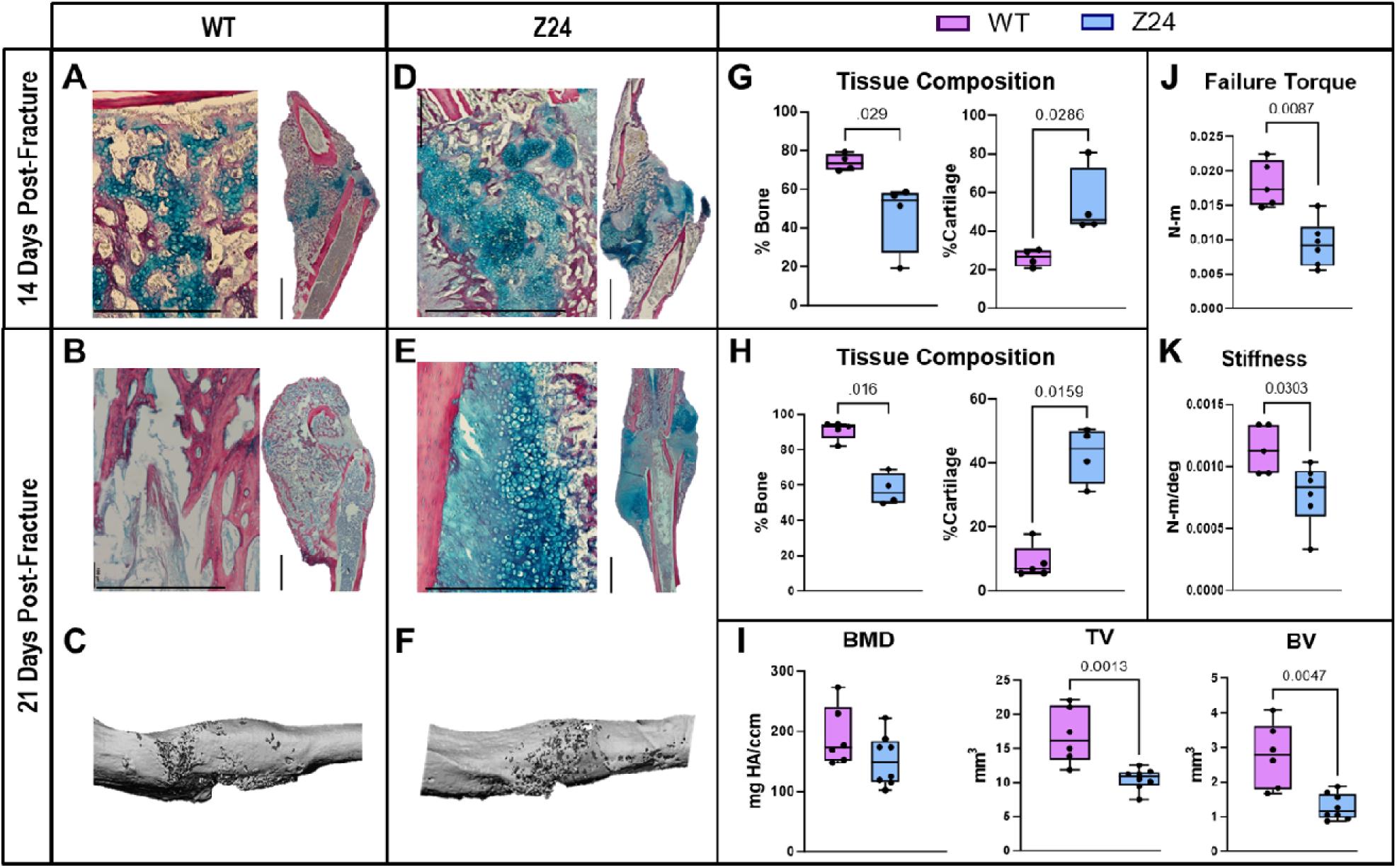
Histomorphometry quantification and. μ**CT analysis reveal Z24^-/-^ mice exhibit decreased torsional bone strength in unfractured tibia and delayed healing after fracture compared to age-matched WT mice**. Hall Brundt’s Quadruple stained images of representative mice tibiae fracture calli **(A, D)** 14 and **(B, E)** 21 days post-fracture (Scale bar: 1 mm (left, 10X), 2 mm (right, 2X)). Histomorphometric quantification of bone and combined cartilage and fibrous tissue volumes in tibial fracture callus **(G)** 14 and **(H)** 21 days after fracture (n: WT = 5, Z24**^-/-^** = 4). Representative μCT images of fracture callus bone architecture at 21 days post-fracture in **(C)** WT and **(F)** Z24**^-/-^** mice. **(I)** Bone mineral density (BMD), total volume (TV), and bone volume (BV) 21 days post-fracture (n: WT = 6, Z24**^-/-^** = 8). **(J)** Maximum torque and **(K)** torsional stiffness in unfractured, contralateral tibia at 21 days post-fracture (n: WT = 5, Z24**^-/-^** = 6). Significance was defined by p ≤ 0.05 determined by Mann-Whitney U test.

### Z24^-/-^ mice have elevated cellular senescence with an increased production of pro-inflammatory cytokines

Flow cytometry analysis of WT and Z24**^-/-^** peripheral blood mononuclear cells (PBMCs) collected pre-fracture demonstrated an average of 76.2% more C_12_FDG^+^ senescent PBMCs in Z24**^-/-^** mice compared to WT mice (p = 0.004) (**Figure 3A-B**). PBMCs isolated 14 days post-fracture demonstrated a 131.9% increase in C_12_FDG^+^ senescence in Z24**^-/-^** mice compared to WT mice (p < 0.001) (**Figure 3C**) with a concomitant systemic increase in the expression of the canonical pro-inflammatory cytokine *Tnf-α* (p = 0.017) (**Figure 3D**). While *Il-1* , another classic pro-inflammatory cytokine, trended higher in Z24**^-/-^** mice compared to WT mice, the difference did not reach significance at the sample size tested (**Figure 3E**).

**Figure 3.**
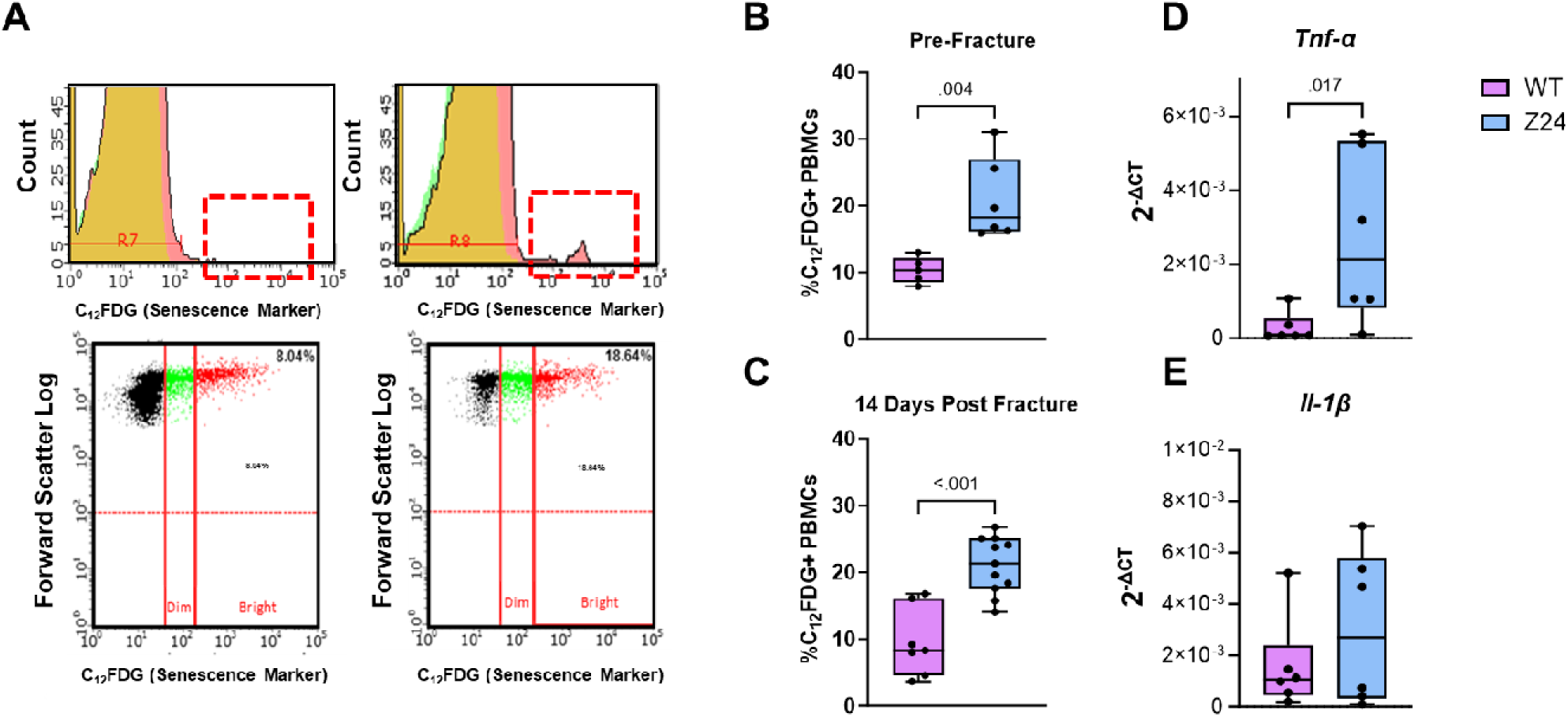
Elevated systemic senescence and SASP expression pre- and post-fracture in Z24^-/-^ mice compared to WT. (A) Representative flow cytometry gating readouts for WT and Z24^-/-^ mice peripheral blood mononuclear cells (PBMCs) stained with C_12_FDG. The C_12_FDG^+^ “bright” cell population (red) is considered senescent. **(B)** C_12_FDG^+^ PBMC senescence prior to fracture (n: WT=5, Z24^-/-^=6) and **(C)** 14 days post-fracture (n: WT=8, Z24^-/-^=12). **(D-E)** Gene expression of inflammatory markers (Tnf- , Il-1 ) in the kidneys 14 days after fracture (n=6). P ≤ 0.05 determined by Mann-Whitney U test.

Multiplex and ELISA analysis of plasma 21 days post-fracture was performed to evaluate systemic differences in a subset of proteins established in the context of aging, senescence, inflammation, and bone healing (IL-13, IP-10, eotaxin, GDF-15, SOST) [8, 35]. Z24^-/-^ mice demonstrated no significant difference in eotaxin, IP-10, and GDF-15 (**Figure 4B-D**). However, there was a 2.4-fold higher level of IL-13 in Z24^-/-^ mice (p = 0.0242, **Figure 4A**) and a significantly lower level of SOST (p = 0.0031, **Figure 4E**).

**Figure 4.**
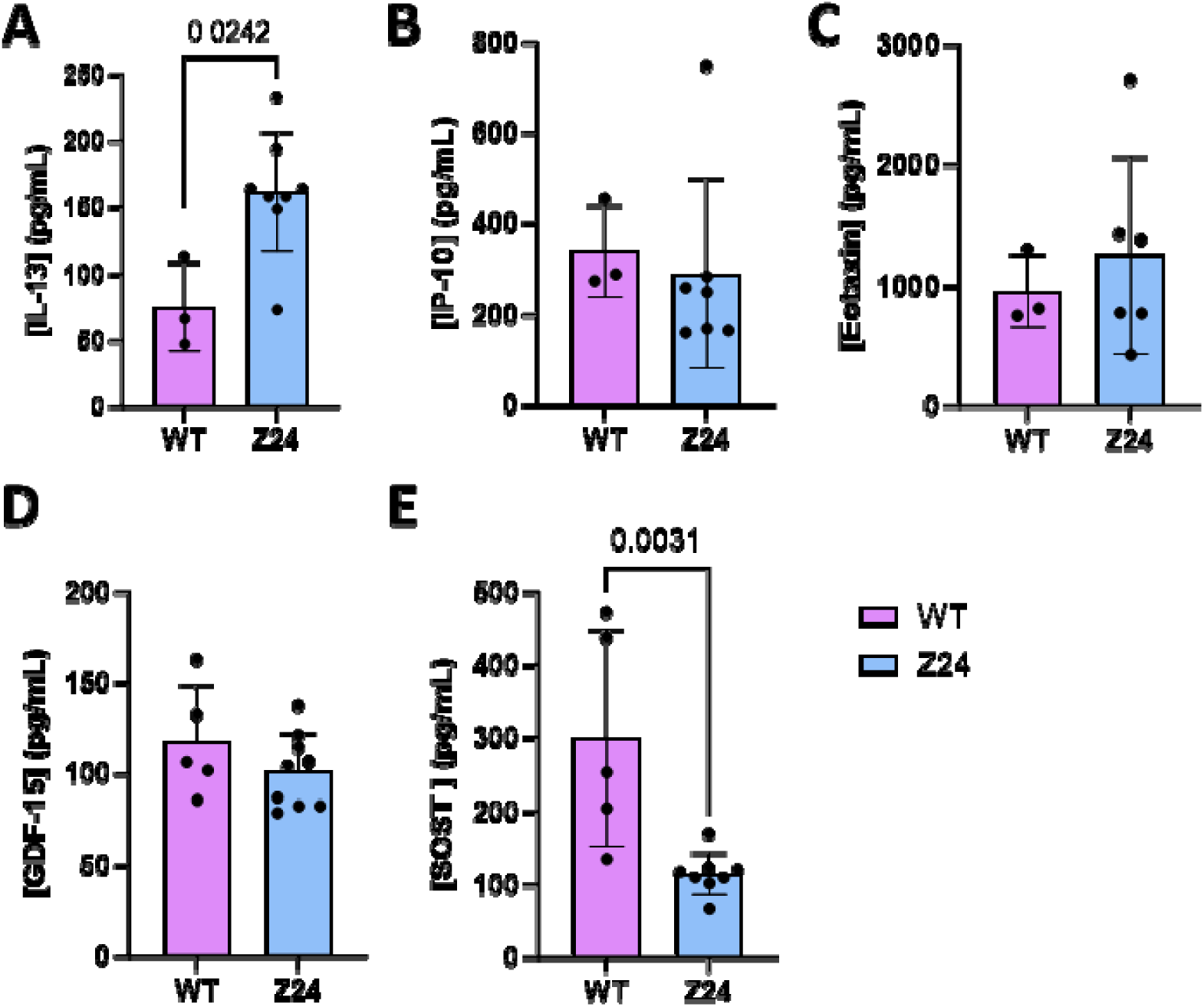
Z24^-/-^ mice display differential plasma protein expression of osteogenic and inflammatory mediators after fracture. Multiplex protein analysis of plasma collected from Z24^-/-^ and WT mice at 21 days post-fracture evaluating the concentration of **(A)** IL-13, **(B)** IP-10, and **(C)** Eotaxin (n: WT = 3, Z24**^-/-^** = 6-8). Quantification via ELISA of plasma proteins collected from Z24^-/-^ and WT mice 21 days post-fracture of **(D)** GDF-15 and **(E)** SOST (n: WT = 5, Z24**^-/-^** = 8-9). Significance was defined as p ≤ 0.05 and tested by Mann-Whitney U test.

### Systemic dysregulation of the innate and adaptive immune system during the time course of fracture repair in Z24^-/-^ mice

Systemic changes to PBMCs collected from peripheral blood were evaluated before injury and at days 3, 14, and 21 post-fracture through precision immunophenotyping to identify all major

myeloid and lymphoid cell types using 30-channel spectral flow. (**Supplemental Table 1-2, Supplemental Figure 1**). At day 3 post-fracture, Z24^-/-^ PBMCs contained significantly fewer CD163^++^ (p=0.0394, **Figure 5A**) and CD206^++^ (p=0.0407, **Figure 5B**) macrophages compared to the age-matched WT mice. No significant differences were found in the relative quantities of neutrophils between the Z24^-/-^ and WT mice (p=0.7044, data not shown).

**Figure 5.**
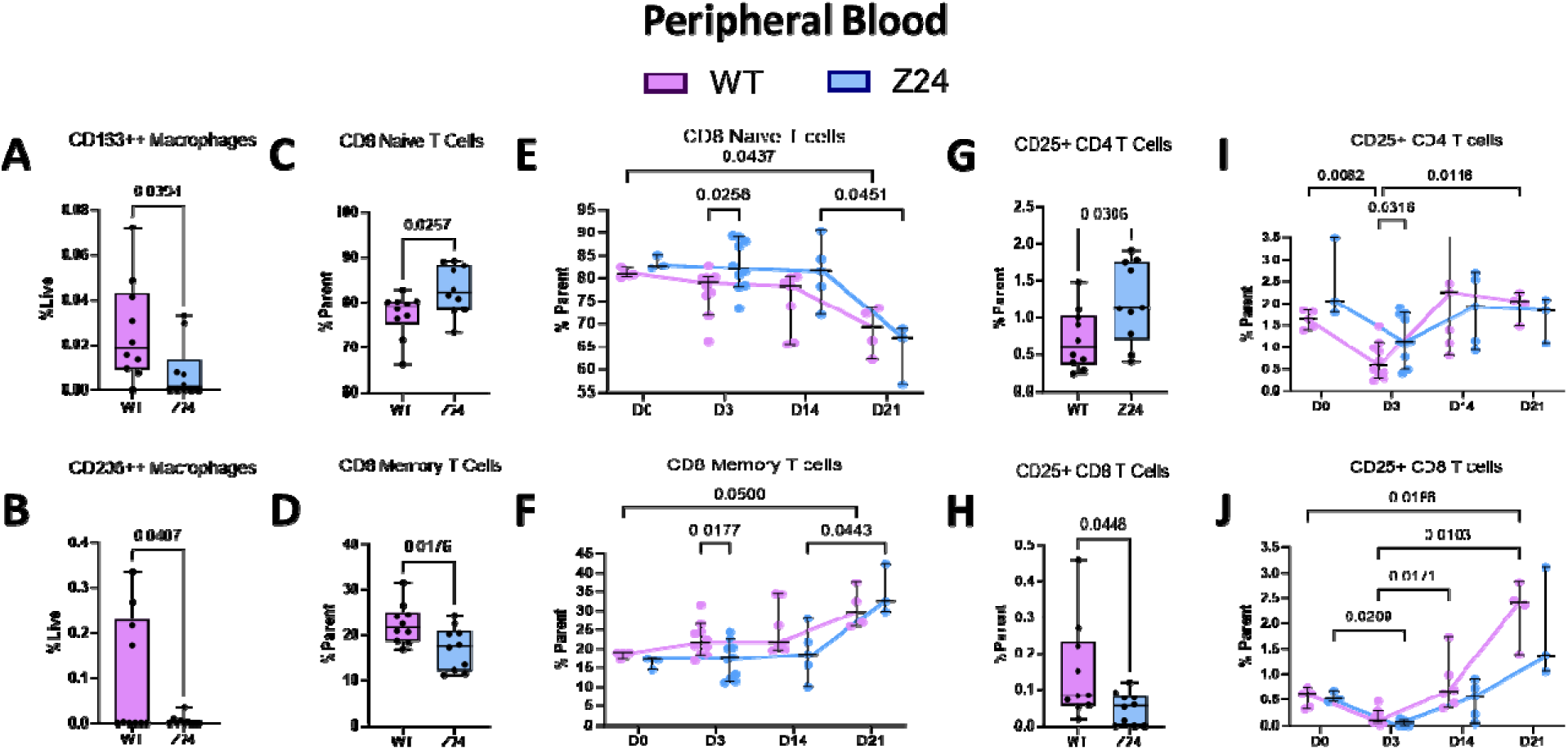
Reduced systemic anti-inflammatory macrophages and unique T cell repertoire in Z24^-/-^. Spectral flow immunophenotyping analysis of blood collected 3 days post fracture evaluating the quantity of **(A)** CD163^++^ macrophages, **(B)** CD206^++^ macrophages, **(C)** CD8^+^ naïve T cells, **(D)** CD8^+^ memory T cells, **(G)** CD25^+^CD4^+^ T cells, and **(H)** CD25^+^CD8^+^ T cells (n:WT=10, Z24^-/-^ = 10). p 0.05 determined by two-tailed T-test. Time course analysis of Z24**^-/-^**blood collected before fr^≤^acture and 3-, 14-, and 21 days post-fracture evaluating the quantity of **(E)** CD8^+^ naïve T cells, **(F)** CD8^+^ memory T cells, **(I)** CD25^+^CD4^+^ T cells, and **(J)** CD25+ CD8^+^ T cells. p<0.05 determined by Mixed-effects analysis (two-way ANOVA) with Tukey multiple comparisons correction test.

Pronounced differences were also displayed in the adaptive immune system phenotype, specifically related to T cell populations. At day 3 following fracture, the Z24^-/-^ mice had a higher proportion of CD8^+^ naïve T cells (p=0.0257, **Figure 5C**) and a smaller proportion of CD8^+^ memory T cells (p=0.0176, **Figure 5D**). Within both the Z24^-/-^ and WT mice peripheral blood, the percentage of naïve CD8^+^ T cells stayed relatively constant throughout the endochondral phase of repair, but then demonstrated a relative decrease at day 21 (p=0.0451, **Figure 5E**). The percentage of CD8^+^ memory T cells similarly stayed constant through day 14, but here increased 21 days post fracture, concomitant with the decrease in naïve CD8^+^ T cells (p=0.0443, **Figure 5F**).

At day 3, we also found a higher number of activated CD25^+^CD4^+^ T helper cells in the Z24^-/-^ mice (p=0.0306, **Figure 5G**) but a smaller proportion of activated CD25^+^CD8^+^ cytotoxic T cells (p=0.0448, **Figure 5H**). While activated CD25^+^CD8^+^ cytotoxic T cells reached their lowest levels in the blood in both mice genotypes at day 3 (p=0.0209), they rose steadily in quantity to significantly accumulate at day 21 (**Figure 5J**). Z24^-/-^ and WT mice displayed similar quantities and subsets of B cells in circulation at all time points (p=0.5119, data not shown).

### Z24^-/-^ mice have increased myelopoiesis and diminished lymphopoiesis three days following fracture

Based on the observed differences in the circulating PBMCs, we next completed detailed immunophenotyping of the ipsilateral (fractured tibia) and contralateral (unfractured tibia) bone marrow compartments. (**Figure 6-7**) While all time points were analyzed (data not shown), the most significant differences were observed during the pro-inflammatory phase of fracture healing 3 days post-fracture. Z24^-/-^ mice displayed significant evidence of diminished lymphopoiesis in the bone marrow on both the ipsilateral and contralateral sides 3-days after fracture. This is evidenced by decreased B cell precursors (p=0.0053, p=0.0051, respectively;**Figure 6A-B**), decreased immature B cells (p=0.0012, p=0.0029, respectively; **Figure 6C-D**), and increased quantity of non-B cells (p=0.0056, p=0.0052, respectively; **Figure 6E-F**).

**Figure 6.**
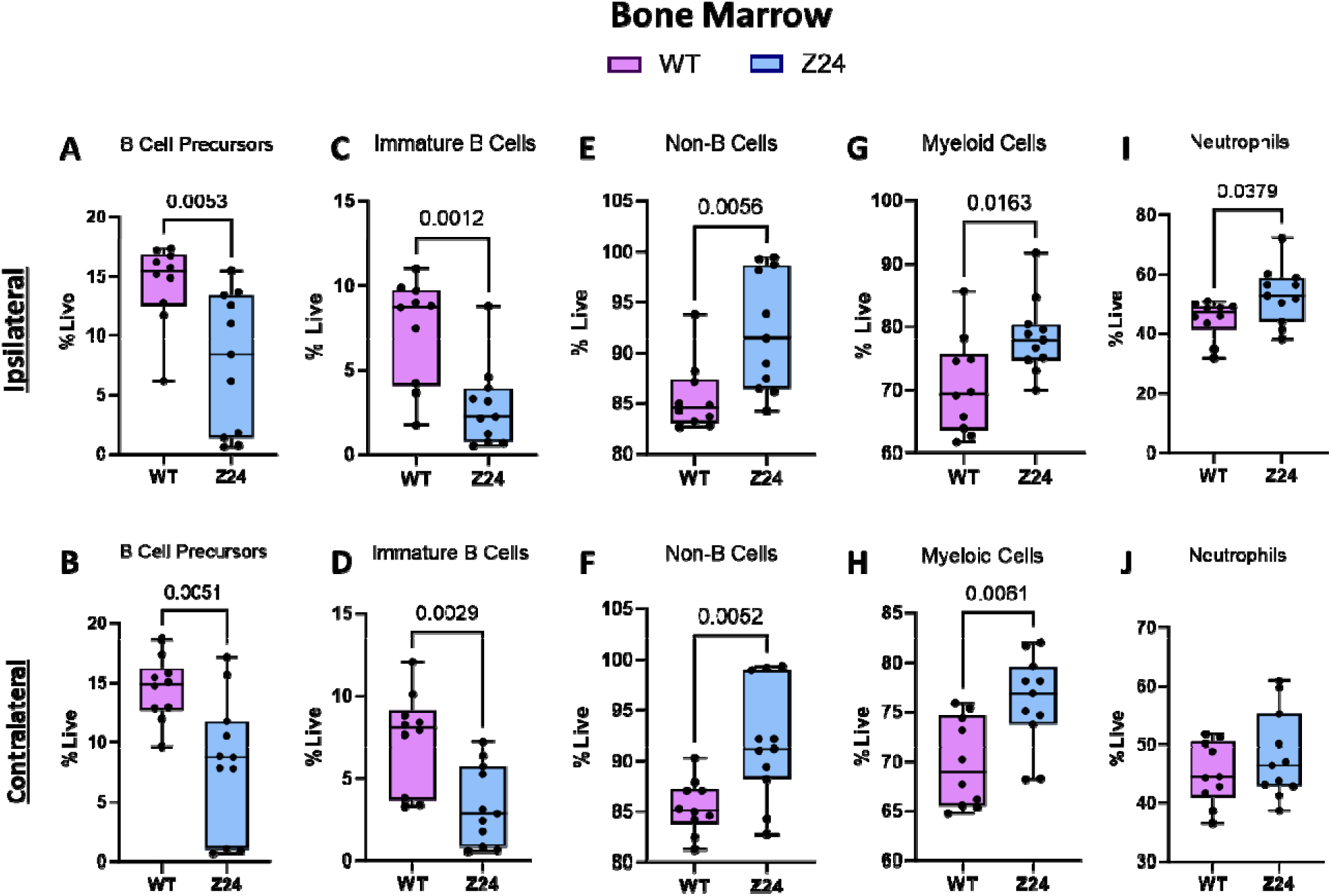
Differential B cell development, myeloid and neutrophil populations in Z24^-/-^ bone marrow 3 days post-fracture. Spectral flow immunophenotyping analysis of ipsilateral and contralateral tibia bone marrow collected 3 days post-fracture evaluating the quantity of live **(A-B)** B cell precursors, **(C-D)** immature B cells, **(E-F)** non-B cells, **(G-H)** myeloid cells, and **(I-J)** neutrophils. (n: WT=10, Z24^-/-^ = 11). P 0.05 determined by two-tailed T-test.

**Figure 7.**
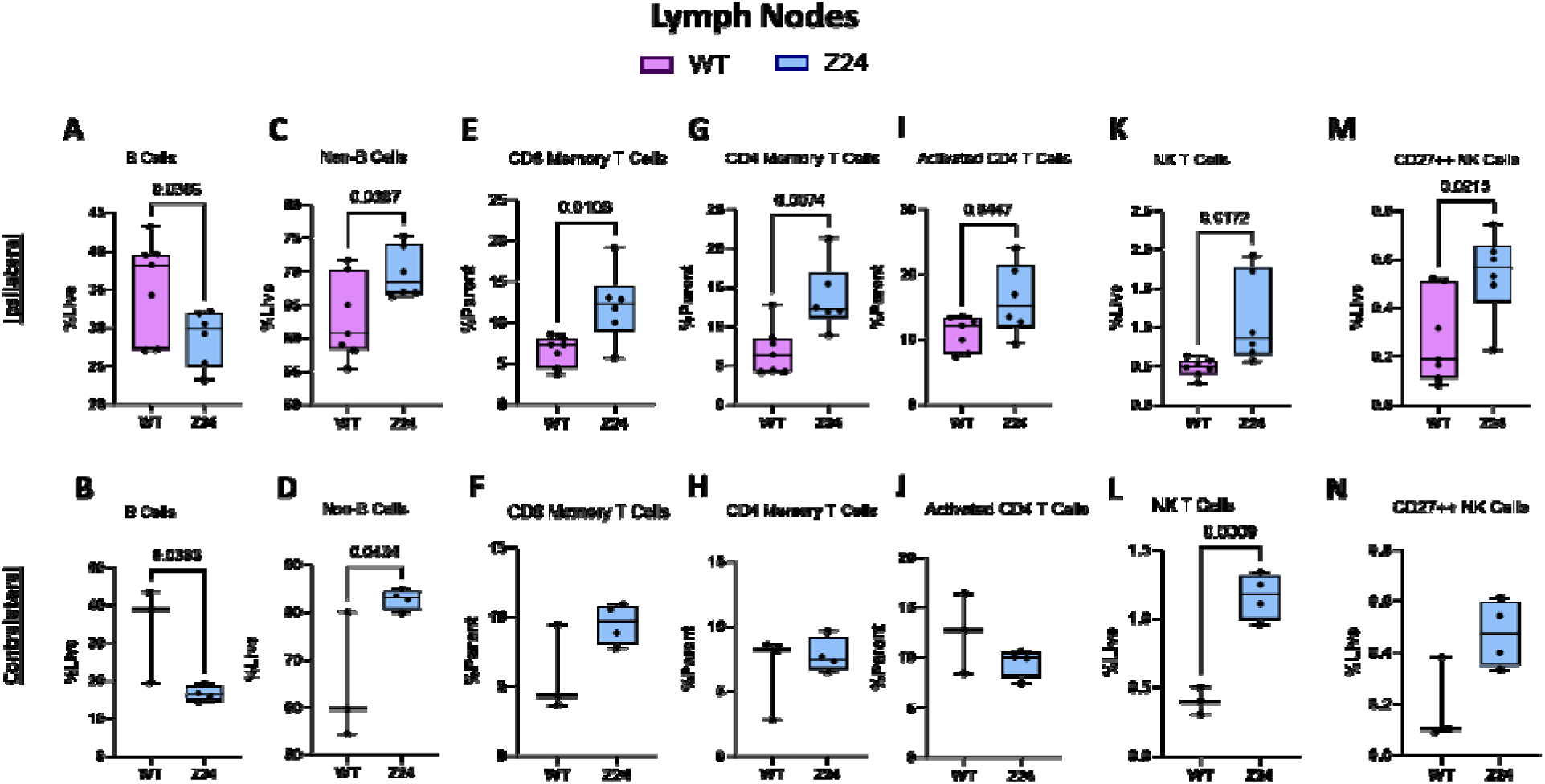
Differential B cell population, T cell activation, and natural killer cell repertoire in Z24^-/-^ ipsilateral lymph nodes. Spectral flow immunophenotyping analysis of ipsilateral and contralateral inguinal lymph nodes collected 3 days post-fracture evaluating the quantity of **(A-B)** B cells, **(C-D)** non-B cells, **(E-F)** CD8^+^ memory T cells, **(G-H)** CD4^+^ memory T cells, **(I-J)** activated CD4^+^ T cells, **(K-L)** NK T cells, and **(M-N)** CD27^++^ NK cells. (n: WT=3-7, Z24**^-/-^** = 4-6). p ≤ 0.05 determined by two-tailed T-test.

Conversely, Z24^-/-^ mice presented with increased myeloid cells in both the ipsilateral (p=0.0163) and contralateral (p=0.0061) limbs (**Figure 6G-H**). Neutrophils were only significantly increased in the ipsilateral limb (p=0.0379, **Figure 6I-J**). The macrophage populations were not significantly different in the bone marrow of the Z24^-/-^ mice compared to WT mice (data not shown).

Z24^-/-^ mice again displayed signs of diminished quantity of B cells on both the ipsilateral and contralateral sides in the inguinal lymph nodes, evidenced by decreased B cell populations (p=0.0386 **Figure 7A**; p=0.0383, **Figure 7B**) and increased quantity of non-B cells (p=0.0387, **Figure 7C**; p=0.0434, **Figure 7D**). There were also significant changes to the T cell populations with increased CD4^+^ (p=0.0074, **Figure 7G**) and CD8^+^ memory T cells (p=0.0106, **Figure 7E**), and activated CD25^+^CD4^+^ T helper cells (p=0.0447, **Figure 7I**) in Z24^-/-^ mice relative to WT mice. This hyperactivated T cell response is only present in the ipsilateral lymph nodes (**Figure 7F, H, J**). Z24^-/-^ mice also presented with significantly increased numbers of natural killer (NK) T cells on both the ipsilateral (p=0.0172, **Figure 7K**) and contralateral (p=0.0009, **Figure 7L**) lymph nodes and increased CD27^++^ NK cells (p=0.0215, **Figure 7G**) in the ipsilateral lymph node.

## DISCUSSION

Several studies have utilized the Z24^-/-^ progeria mouse to model aging-associated musculoskeletal conditions and frailty [25, 26]. Most relevant to our work, multiple studies have established that by three months of age Z24^-/-^ mice present with an osteoporotic/osteopenic phenotype with decreased bone mineral density, loss of trabecular and cortical bone, and increased trabecular spacing, along with reports of lower weight, loss of hair, and a malnourished appearance [36–39]. This accelerated aging phenotype is caused by a mutation in the Z24 metalloproteinase that leads to nuclear blebbing and cellular stiffness due to the disrupted assembly of lamin in the cytoskeleton, a process that also occurs during natural aging [36, 40–43]. It has also been demonstrated that reduced lamin A/C [44] can lead to impaired osteogenesis with increased adipogenesis [45–47], and that frail individuals have reduced circulating osteoprogenitors [28].

Our research is the first to explore the Z24^-/-^ mouse model in the context of fracture healing. General skeletal frailty in Z24^-/-^ mice at the time of fracture (10-14 weeks of age) was confirmed through the decreased failure torque and torsional stiffness in the tibae of the Z24^-/-^ mice. Consistent with murine experiments demonstrating delayed fracture healing in naturally aged mice [13, 14, 48], Z24^-/-^ mice presented with retained cartilage and decreased bone fractions in the fracture callus compared to age-matched young WT mice. The Z24^-/-^ mice also developed a

smaller overall fracture callus, similar to other murine models of impaired fracture healing [17,

18, 49]. As previously published, this is most likely due to the depletion of the functional periosteal stem cell progenitor population [13, 49].

To understand the underlying phenotype that may contribute to delayed fracture repair in Z24^-/-^ mice, we measured the senescent and immunophenotype of the progeric, Z24^-/-^ mice relative to age-matched WT counterparts. The most widely used biomarker of cellular senescence is increased activity of the acidic senescence-associated β β study, we quantified systemic senescence using C_12_FDG flow cytometry of the peripheral blood mononuclear cells (PBMCs). C_12_FDG is a compound that fluoresces at 514 nm when hydrolyzed by intracellular SA-β-gal and has been well described for use in mice [32, 33] and, β more recently, our work in humans [51]. Using this technique, we found that Z24^-/-^ progeria mice had a significantly increased number of C_12_FDG^+^ PBMCs before fracture compared to the age-matched WT mice, with an even further relative accumulation of these cells 14-days following fracture.

Immune dysregulation with natural aging, commonly referred to as “inflammaging” [52], is causal in age-related morbidities of solid organs [53, 54] and delayed fracture repair [14, 48]. We found the increased number of circulating C_12_FDG^+^ senescent PBMCs in Z24^-/-^ mice was associated with a systemic pro-inflammatory state characterized by upregulated *Tnf-α* during the endochondral phase of healing. A pro-inflammatory state, defined by increased systemic *Tnf-α*, has also been described in murine models of delayed fracture healing due to natural aging [48, 55], diabetes [17], and rheumatoid arthritis (RA) [18]. We also analyzed a limited number of candidate cytokines for differential systemic expression 21 days post-fracture based on prior work suggesting predictive relationships with bone regeneration after trauma [35]. In our model, we found very few of these cytokines had differences in expression, with the exception of IL-13, which was significantly upregulated in Z24^-/-^ mice fracture compared to the WT mice. IL-13 has a high degree of structural homology with IL-4 and is commonly considered an anti-inflammatory cytokine. In fracture healing, early upregulation of IL-13 was recently identified as a biomarker of successful healing in femoral defects with delayed treatment in otherwise healthy young adult-age rats [35]. However, IL-13 is also associated with chronic inflammatory diseases leading to fibrosis, systemic sclerosis, and asthma. [56, 57] IL-13 is also made by NK and activated T helper cells, both of which were found to be higher in the draining lymph node, but not the contralateral lymph node, of Z24^-/-^ mice compared to WT [57]. Consequently, in this Z24^-/-^ model of accelerated aging, the increased IL-13 at the late stages of healing where we measured expression may instead be associated with a dysregulated immune state and contribute to the increased fibrosis observed in the fracture callus of the Z24^-/-^ mice.

To characterize differences in the immunophenotype between the Z24^-/-^ mice and age-matched WT mice, we completed spectral flow cytometry on bone marrow, lymph nodes, and peripheral blood to identify a broad spectrum of the major myeloid and lymphoid cell populations. The systemic immune response was analyzed by immunophenotyping of the peripheral blood and compared to the local immune response within the bone marrow and inguinal lymph node that drains the tibial fracture site. Significant differences in both the innate and adaptive immune systems were observed between the two mice cohorts. Within the innate response, 3 days after fracture, we saw significantly more neutrophils and myeloid cells within the bone marrow of Z24^-/-^ mice and fewer circulating anti-inflammatory macrophages. Neutrophils are the most abundant cells recruited to the fracture site in the inflammatory stage and release damage associated molecular patterns (DAMPs) that are critical to initiating the endogenous innate immune response but are associated with poor outcomes when present in excess. Supporting this, preclinical neutrophil depletion studies demonstrate they are essential to the normal healing process [58], but most clinical studies show that higher neutrophil concentrations are associated with an increased risk of severe complications, delayed union, and even mortality in geriatric patients (as recently reviewed [59]). These prior studies suggest that the higher concentration of neutrophils in the Z24^-/-^ mice may contribute to the delayed healing phenotype that we observed.

The increased concentration of neutrophils was concomitant with an increase in myeloid cells and a decrease in B lymphocytes, which are hallmark immune changes observed with natural aging. [60, 61] While we did see a higher number of total myeloid cells in the bone marrow, we did not see an increase in the local pro-inflammatory macrophages. Rather, we measured a significant reduction in the number of anti-inflammatory CD163^+^ and CD206^+^ circulating macrophages 3 days following fracture, with these lower quantities generally being maintained throughout the time course of healing. While we did not measure the pro-inflammatory macrophages directly within the fracture callus, these results indicate either that Z24^-/-^ mice do not have the same upregulation of macrophages noted in naturally aged mice [13, 14], or that the Z24^-/-^ mice fail to mount the necessary accumulation of pro-inflammatory macrophages required to initiate healing [62, 63].

Significant adaptive immune system changes were also observed systemically and locally in Z24^-/-^ mice relative to age-matched WT mice. In the bone marrow and draining lymph nodes the, Z24^-/-^ mice demonstrated a dramatic reduction in the number of B cells. This finding is consistent with literature showing that a reduction in lymphopoiesis is a hallmark of aging [60]. Moreover, an infiltration of B cells is one of the strongest positive predictors of optimal bone healing [35], supporting our correlation between the poor fracture healing in the Z24^-/-^ mice and decreased B-cells in the draining lymph node.

The T cell phenotype in the Z24^-/-^ mice was more difficult to equate to existing literature due to the diversity in T cell labeling strategy used across the various flow cytometry protocols. In our study we found Z24^-/-^ mice had higher numbers of naïve CD8^+^ T cells and reduced differentiated cytotoxic CD8^+^ T cells (both memory and activated) in the blood but an accumulation of cytotoxic cells within the lymph node. We also found higher number of CD4^+^ T-helper cells systemically in the blood and within the lymph node. Notably, the increase in Z24^-/-^ CD4^+^ and CD8+ memory cells was only present in the fracture side (ipsilateral) lymph node, not in the contralateral lymph node or systemically in the blood. This, combined with the increase in Z24^-/-^ myeloid cells found in the bone marrow, suggests that elevated myeloid abundance and activation may promote a hyperactivation of T cells local to the fracture side. This Increased local CD4^+^ T-helper and cytotoxic CD8^+^ T cells was also observed in middle-aged mice with a similar reduction in mobilization of the cytotoxic CD8^+^ T cells, but in this study they did not find increased systemic T-helper cells in the middle-aged mice [64]. Elevated cytotoxic T cells within the fracture callus was also seen in aged mice in the recently published Molitoris *et al* study, however this result did not reach significance, perhaps due to a relatively low sample size [55]. Further, previously published work has found that an increase in CD8^+^ effector memory T cells (CD3^+^CD8^+^CD11a^++^CD28^−^ CD57^+^) was highly correlated to delayed fracture healing [65]. Taken together, the T cell dynamics of the progeria mice relative to age-matched WT mice share similarities with published work looking at the difference between young and aged mice and those with delayed fracture healing.

Lastly, we also found a significant increase in CD27^++^ NK and NK T cells in the Z24^-/-^ mice. To our knowledge, there is no comparative research currently available on these cell types during fracture healing, and therefore, these cells represent an interesting avenue for future research. In general, the role of NK cells in trauma is not well understood, and a recent review of the literature suggests that there is a tight interplay between NK cells and mesenchymal stromal cells during fracture healing [59].

This study has some noted limitations. The pathology of bone fracture healing is complex and the different phases of fracture repair present shifting biological environments that may not have been equivalent in the two mice populations due to the “accelerated” timescale of aging of the Z24^-/-^ mice. Furthermore, the process of aging in general is not universal even in congenic, co-housed mice, evidenced by some elderly animals presenting as “super-agers” with immunological states that align more closely with young mice [14]. Similarly, we found more variance in the quantitative fracture healing and immunosenescent outcomes of Z24^-/-^ mice compared to the age-matched WT mice. Although we did not find any “super-ager” outliers in the cohort tested, we did see gross phenotypic differences across Z24^-/-^ mice. We also acknowledge that due to the limitation of Mendelian frequency of the Z24^-/-^ mice from the Z24^+/-^ parents, we were unable to segregate our data set by mouse sex. As such, significant findings could be more profound through the future segregation of sex, since sex-differences have recently been demonstrated in naturally aged mice [55]. Lastly, the Z24^-/-^ mice are smaller than their age-matched WT counterparts with a frail behavioral phenotype that we, and others, have also shown are associated with osteoporotic and sarcopenic standards. This is in contrast to naturally aged mice in controlled breeding settings which tend to be much larger than young adult mice; however, frailty, sarcopenia, and osteoporosis are characteristics of aged humans. These different weight spectrums between Z24^-/-^, young adult WT, and naturally aged WT mice may also influence healing outcomes as there is an interconnected nature between bone loading and fracture repair.

Despite these limitations, here we present Z24^-/-^ mice as a practical murine model of delayed healing that has many parallels to aging associated co-morbidities that drive delayed fracture healing in human populations. There was previously a gap in models that enabled testing of therapeutic approaches to accelerate fracture healing in situations of delayed bone repair that translates well to clinical experiences. The increased senescent burden and age related changes to lymphopoiesis and increased neutrophil infiltration lend themselves nicely to testing of senolytics during fracture healing which is emerging as an exciting new approach to address age-related delays in bone healing [66].

## SUPPLEMENTARY

Supplementary Figure 1. Immunophenotyping Gating Strategy Example A

## AUTHOR CONTRIBUTIONS

**Victoria R. Duke:** Formal Analysis, Investigation, Data Curation, Writing - Original Draft, Writing- Review & Editing, Visualization. **Marc J. Philippon Jr.:** Formal Analysis, Investigation, Data Curation, Writing - Original Draft, Writing - Review & Editing, Visualization. **Dane R.G. Lind:** Formal Analysis, Investigation, Data Curation, Writing - Original Draft, Writing- Review & Editing, Visualization. **Herbert Kasler:** Methodology, Formal Analysis, Investigation, Data Curation, Resources, Writing – Review and Editing. **Kohei Yamaura:** Investigation. **Matt Huard:** Investigation**. Molly Czachor:** Investigation, Writing – Review and Editing. **Justin Hollenbeck:** Investigation. **Justin Brown:** Investigation. **Alex Garcia:** Investigation. **Naomasa Fukase:** Investigation. **Ralph S. Marcucio:** Methodology, Validation, Writing – Review and Editing. **William S. Hambright:** Conceptualization, Methodology, Validation, Supervision, Writing – Review and Editing. **Anna-Laura Nelson:** Investigation, Writing – Review and Editing**. Dustin** 1. **M. Snapper:** Formal Analysis, Investigation, Writing - Original Draft, Writing - Review & Editing.**Johnny Huard:** Conceptualization, Methodology, Resources, Funding Acquisition, Writing – Review and Editing. **Chelsea S. Bahney:** Conceptualization, Methodology, Resources, Funding Acquisition, Supervision, Formal Analysis, Writing – Original Draft, Writing - Review & Editing.

## ACKNOWLEDGEMENTS

The authors thank Laura Chubb and the Lab Animal Resources Staff at Colorado State University for their help with the animal surgeries and μCT analysis, and Suzanne Page for grant management and operations at the Steadman Philippon Research Institute.

---

**Supplementary Figure 1.**
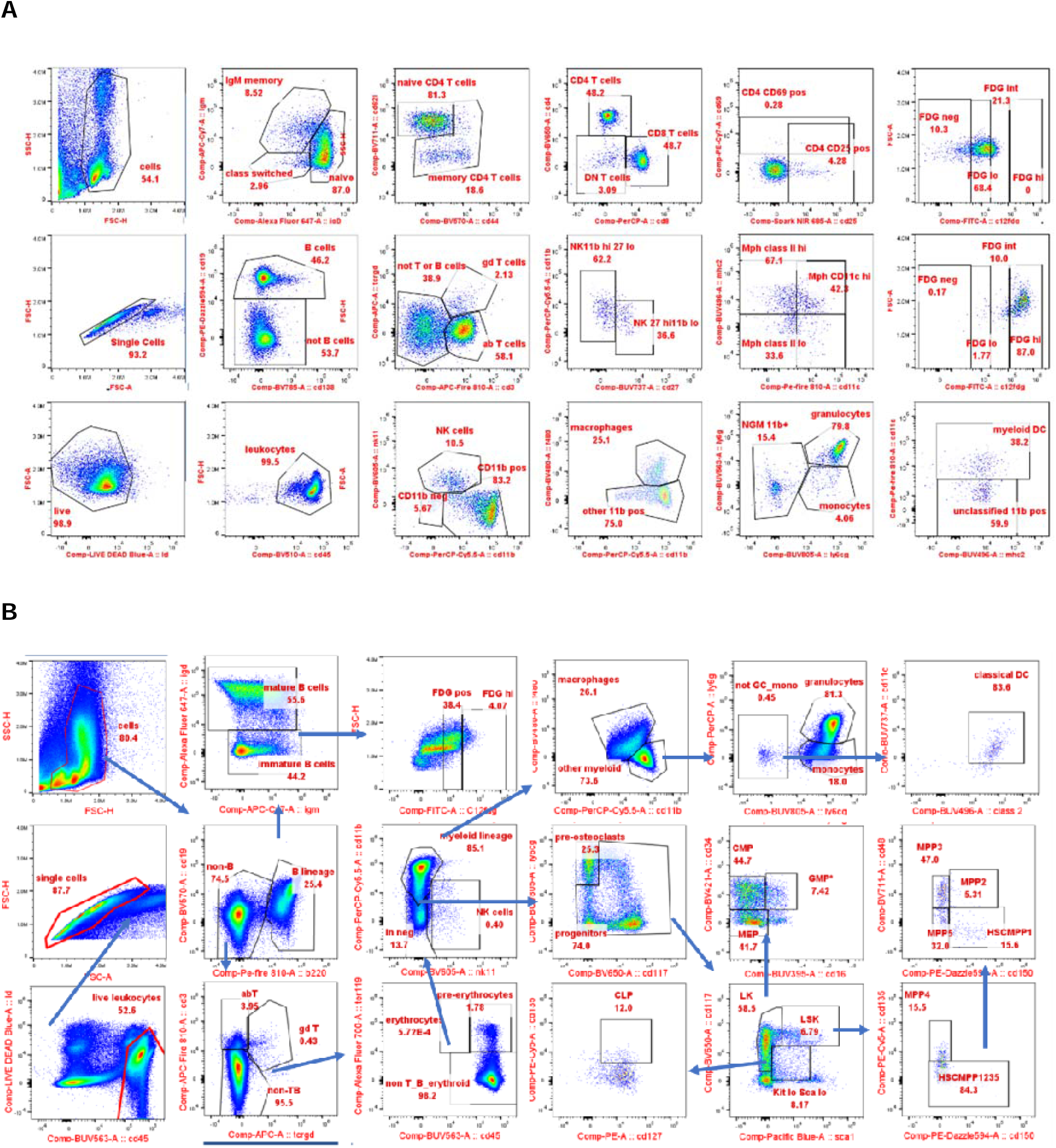
Immunophenotyping gating strategy. (A) Representative gating with 30-color panel for main populations in blood and tissues. (B) Representative gating with 30- color panel for main populations in bone marrow.

**Supplementary Table 1.**
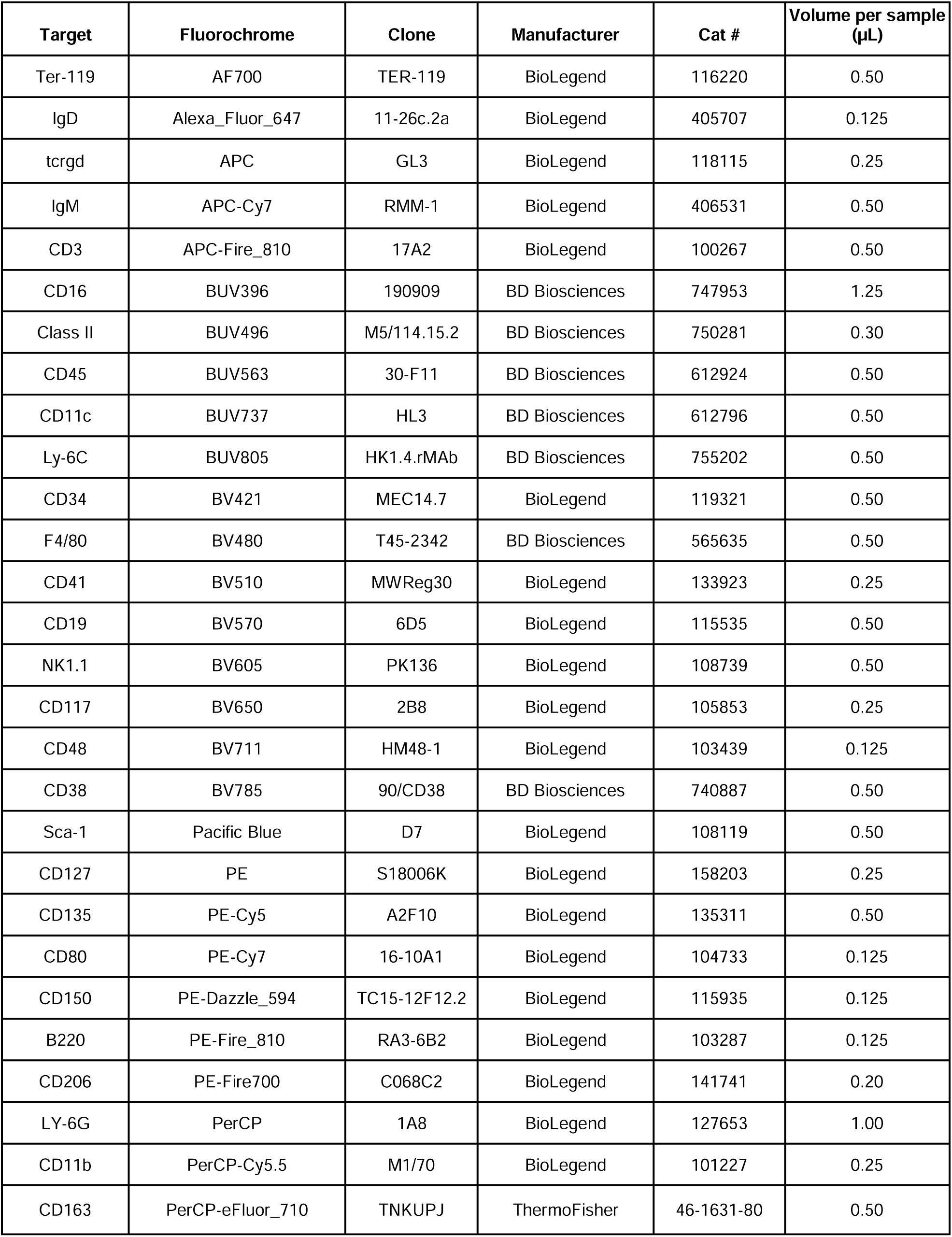
Antibodies for immunophenotyping.

**Supplementary Table 2.**
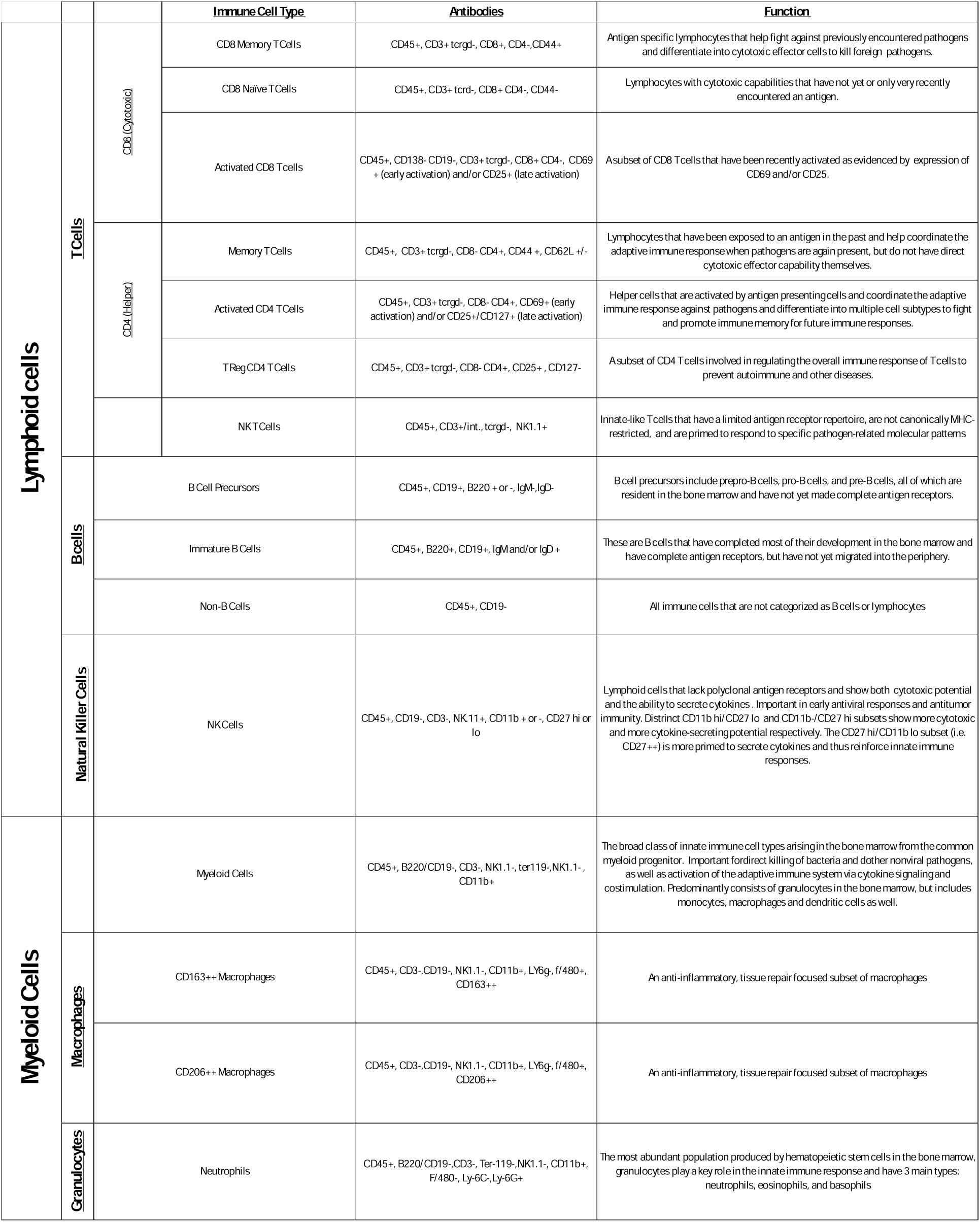

